# Prostaglandin D_2_ Signaling is Not Involved in the Recovery of Rat Hindlimb Tendons from Injury

**DOI:** 10.1101/778126

**Authors:** Dylan C Sarver, Kristoffer B Sugg, Jeffrey R Talarek, Jacob B Swanson, David J Oliver, Aaron C Hinken, Henning F Kramer, Christopher L Mendias

## Abstract

Injured tendons heal through the formation of a fibrovascular scar that has inferior mechanical properties compared to native tendon tissue. Reducing inflammation that occurs as a result of the injury could limit scar formation and improve functional recovery of tendons. Prostaglandin D_2_ (PGD_2_) plays an important role in promoting inflammation in some injury responses and chronic disease processes, and the inhibition of PGD_2_ has improved healing and reduced disease burden in animal models and early clinical trials. Based on these findings, we sought to determine the role of PGD_2_ signaling in the healing of injured tendon tissue. We tested the hypothesis that a potent and specific inhibitor of hematopoietic PGD synthase (HPGDS), GSK2894631A, would improve the recovery of tendons of adult male rats following an acute tenotomy and repair. To test this hypothesis, we performed a full-thickness plantaris tendon tenotomy followed by immediate repair and treated rats twice daily with either 0mg/kg, 2mg/kg, or 6mg/kg of GSK2894631A. Tendons were collected either 7 or 21 days after surgical repair, and mechanical properties of tendons were assessed along with RNA sequencing and histology. While there were some differences in gene expression across groups, the targeted inhibition of HPGDS did not impact the functional repair of tendons after injury as HPGDS expression was surprisingly low in injured tendons. These results indicate that PGD_2_ signaling does not appear to be important in modulating the repair of injured tendon tissue.

## Introduction

Tendon is a dynamic tissue that is important for transmitting and storing elastic energy between skeletal muscle and bone. While tendon is mechanically robust, it can rupture in response to excessive strain placed on the tissue, or with repetitive high frequency loading activities that generate a series of small tears which propagate over time (Sharma & Maffulli, 2006; Mead *et al.*, 2018). Tendon ruptures can be treated either conservatively or with surgical repair, but in both cases a fibrovascular scar forms between the torn tendon stumps (Sharma & Maffulli, 2006; Yang *et al.*, 2013; Ganestam *et al.*, 2016). This scar tissue has inferior mechanical properties compared to native tendon tissue and disrupts the normally efficient transfer of force throughout the tendon, which leads to impaired locomotion (Yang *et al.*, 2013; Nourissat *et al.*, 2015; Freedman *et al.*, 2017).

There is a substantial inflammatory response that occurs in the early stages of the repair of a torn tendon, including infiltration of neutrophils and macrophages, and an upregulation in proinflammatory cytokines and cyclooxygenase (COX) enzymes (Marsolais *et al.*, 2001; Koshima *et al.*, 2007). Nonsteroidal antiinflammatory drugs (NSAIDs) and COX-2 inhibitors (coxibs) have been used clinically to treat pain and prevent inflammation after tendon repair, but in most cases the use of NSAIDs or coxibs reduces or delays tissue healing (Ferry *et al.*, 2007; Dimmen *et al.*, 2009; Hammerman *et al.*, 2015). This is true not only for tendon, but also for other musculoskeletal tissues including skeletal muscle, bone, and the enthesis (Cohen *et al.*, 2006; Su & O’Connor, 2013; Dueweke *et al.*, 2017; Lisowska *et al.*, 2018). NSAIDs and coxibs block the production of prostaglandin H_2_ (PGH_2_) from arachidonic acid, and PGH_2_ is a precursor for the production of several prostaglandins including PGD_2_, PGE_2_, PGF_2α_, and PGI_2_ (Trappe & Liu, 2013). Although less is known for tendon, the negative effects of NSAIDs and coxibs on skeletal muscle healing are thought to occur by blocking the production of PGF_2α_, which is critical for muscle fiber growth and regeneration (Trappe & Liu, 2013). Therefore developing a therapy that can specifically target proinflammatory prostaglandins without impacting other prostaglandins could improve the treatment of tendon disorders.

PGD_2_ is a proinflammatory prostaglandin that is produced from PGH_2_ by two enzymes, hematopoietic PGD synthase (HPGDS) and lipocalin-type PGD synthase (PTGDS) (Joo & Sadikot, 2012; Thurairatnam, 2012). HPGDS is expressed in various immune and inflammatory cells that participate in the repair of injured tissues (Thurairatnam, 2012), and the targeted inhibition of PGD_2_ production improves skeletal muscle repair after injury and also reduces the pathological muscle changes in the *mdx* model of Duchenne muscular dystrophy (Mohri *et al.*, 2009; Thurairatnam, 2012). Blocking PGD_2_ production has also improved outcomes in animal models and small clinical trials of pulmonary, autoimmune, and neurodegenerative disease, among others (Thurairatnam, 2012). Based on these findings, we sought to test the hypothesis that the targeted inhibition of PGD_2_ would improve tendon healing following a plantaris tenotomy and repair. To test this hypothesis, we induced an acute plantaris tendon tear followed by an immediate repair, and then treated rats twice daily with GSK2894631A to inhibit the enzymatic activity of HPGDS. Tendons were collected either 7 or 21 days after surgical repair, and mechanical properties were assessed along with transcriptional and histological measurements to determine the impact of HPGDS inhibition on tendon structure and function after tenotomy and repair.

## Materials and Methods

### Animals

This study was approved by the University of Michigan IACUC (protocol PRO00006079). Three-month old male Sprague Dawley rats were purchased from Charles River (Wilmington, MA, USA) and housed under specific pathogen free conditions. Animals were provided food and water *ad libidum*. There were six experimental groups in the study, with N=12 rats per group, for a total of 72 surgical rats. An additional 5 control rats who did not undergo tenotomy surgery or receive the test compound were used in the study to obtain reference values for assays. We estimated sample size the study based on energy absorption values from a previous study (Mendias *et al.*, 2015b). To detect a 30% difference in energy absorption between vehicle and 6mg/kg doses at the 7 day and 21 day time points, using a power of 80% and an α adjusted from 0.05 for multiple observations, required N=9 per each group. We added 3 additional rats to account for unanticipated losses.

### Surgical Procedure and Administration of Test Compound

Animals were deeply anesthetized with 2% isoflurane, and the skin overlying the surgical site was shaved and scrubbed with 4% chlorhexidine. The animals received a subcutaneous injection of buprenorphine (0.05mg/kg, Reckitt Benckiser, Richmond, VA, USA) for pre-operative analgesia. A longitudinal incision was then performed within the interval between the Achilles and plantaris tendons on each hindlimb. The skin and paratenon were split and retracted to achieve optimal visualization of the plantaris tendon, which is located medial and deep to the Achilles tendon. A full-thickness tenotomy was created in the mid-substance of the plantaris tendon, followed by immediate repair using a Bunnell technique with Ethibond (5-0, Ethicon, Sommerville, NJ, USA). The Achilles tendon was left intact to function as a stress shield for the repaired plantaris tendon. A splash block of 0.2mL of 0.5% bupivacaine was administered, the paratenon was then loosely reapproximated using Vicryl suture (4-0, Ethicon), and the skin was closed with GLUture (Abbott, Abbot Park, IL, USA). After recovery, *ad libitum* weightbearing and cage activity were allowed, and the animals received a second injection of buprenorphine (0.05mg/kg) 12 hours after surgery.

GSK2894631A (7-(Difluoromethoxy)-N-((trans)-4-(2-hydroxypropan-2-yl)cyclohexyl)quinoline-3-carboxamide) which is a potent and specific inhibitor of HPGDS (Deaton *et al.*, 2019), was synthesized and prepared by GlaxoSmithKline (King of Prussia, PA, USA). GSK2894631A was suspended in 0.5% hydroxypropyl methylcellulose:0.1% Tween80 and delivered to rats via oral gavage twice daily at doses of 0mg/kg, 2mg/kg or 6mg/kg. Compounds were provided by GlaxoSmithKline to investigators in a blinded fashion, and identified using a single letter code.

Either 7 or 21 days after the tenotomy and repair surgery, animals were deeply anesthetized with an intraperitoneal injection of sodium pentobarbital (50mg/kg, Vortech Pharmaceuticals, Dearborn, MI, USA). The left plantaris tendon, which was used for mechanical properties testing and histology, was removed by making a full-thickness incision proximal to the myotendinous junction and distal to the calcaneus, in order to preserve the myotendinous junction and enthesis. The left plantaris tendon was then wrapped in saline soaked gauze, and stored at −20°C until use. The right plantaris tendon, which was used for RNA analysis, was removed by making an incision just distal to the myotendinous junction and just proximal to the calcaneus to avoid contaminating muscle or bone tissue, snap frozen in liquid nitrogen, and stored at −80°C until use. Following removal of tendons, animals were humanely euthanized by overdose of sodium pentobarbital and induction of a bilateral pneumothorax.

### Mechanical Properties Measurements

Mechanical properties were measured as modified from previous studies (Mendias *et al.*, 2015a; Sarver *et al.*, 2017). Prior to mechanical tests, tendons were thawed at room temperature and then placed in dish containing PBS. Braided silk suture (4-0, Ashaway Line & Twine, Ashaway, RI, USA) was attached to proximal and distal ends of the tendon using a series of square knots to allow the tendon to be attached to pins for geometric measurements, and to the mechanical properties testing apparatus, without damaging the tendon tissue. The tendon was then transferred to a custom device to measure cross-sectional area (Figure 1A). The device consisted of a trough filled with PBS that contained a sedimentary layer of SYLGARD 184 (Dow Chemical, Auburn, MI, USA) to allow the placement of minutien pins, to which the sutured tendon was attached. The trough was also flanked by prisms that allow for visualization of the side view of the tendon. The tendon was held at just taught length, and CSA was calculated from 5 evenly spaced width and depth measurements from high-resolution digital photographs of both top and side views of the tendon. These measurements were then fit to an ellipse, and the average ellipse area was used as the tendon CSA for mechanical properties measurements.

**Figure 1.**
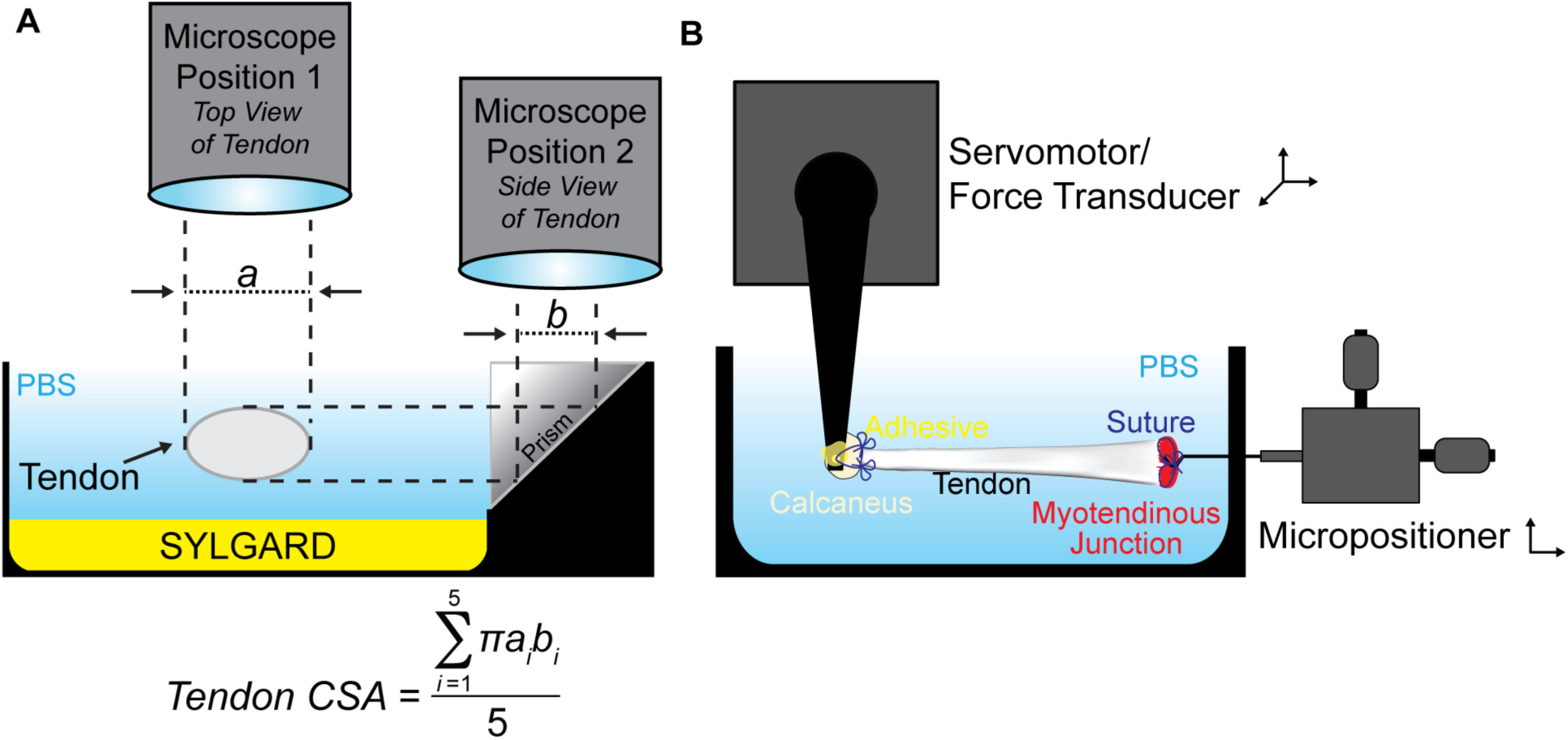
Overview of cross-sectional area and mechanical properties testing devices. (A) Schematic showing the measurement of nominal tendon cross-sectional area, with the tendon shown in cross-section. (B) Schematic showing the measurement of mechanical properties of tendons.

To test mechanical properties, the tendon was then transferred to a bath containing PBS maintained at 25°C. Using the attached sutures, the distal end of the tendon was secured by affixing the calcaneus to a 10N dual-mode servomotor/force transducer (model 305LR, Aurora Scientific, Aurora, ON, Canada), while the proximal end of the tendon was secured at the myotendinous junction to a hook attached to a micropositioner (Figure 1B). Once secured, the tendon was briefly raised up from the bath so that GLUture adhesive could be applied to reinforce the attachment of the calcaneus to the hook. The tendon was then returned to the bath, and its length was adjusted to an approximate 5mN preload, which was consistent with the just taught length, and recorded as L_o_. Each tendon was subjected to 10 load-unload stretch cycles at a constant velocity of 0.05 L_o_/s, and a length change that was 10% of L_o_. Data was recorded using custom LabVIEW software (National Instruments, Austin, TX, USA). Load, stress, tangent modulus, and energy loss were determined for each load-unload cycle. Tangent modulus was defined as the maximum derivative over a 10ms window of data from the stress-strain curve. Energy loss was calculated as the area under force-displacement curve from 10% to 0% strain, subtracted from the area under the force-displacement curve from 0% to 10% strain. Energy loss was then normalized by tendon mass, which was determined by multiplying the volume of tendon by 1.12g/cm^3^ (Ker, 1981).

Following the completion of mechanical properties testing, the tendon ends were trimmed, the tendon was placed in Tissue-Tek OCT Compound (Sakura Finetek, Torrance, CA, USA), flash frozen in isopentane cooled in liquid nitrogen, and then stored at −80°C until use.

### Histology

Longitudinal sections of tendons, approximately 10μm in thickness, were obtained using a cryostat. Sections were stained with hematoxylin and eosin, and digital images were obtained with a Nikon Eclipse microscope equipped with a high-resolution camera (Nikon, Melville, NY, USA).

### RNA Sequencing and Gene Expression

RNA was extracted as modified from previous studies (Nielsen *et al.*, 2014; Gumucio *et al.*, 2014). Tendons were finely minced, and then placed into 2mL tubes containing 2.3mm steel beads and TRI Reagent (Molecular Research Center, Cincinnati, OH, USA), homogenized for 15 sec, and isolated following product directions. The subsequent RNA pellet was then further cleaned up using miRNeasy kit (Qiagen, Valencia, CA, USA), supplemented with DNase I (Qiagen). RNA concentration was determined using a NanoDrop (ThermoFisher Scientific, Waltham, MA, USA), and quality was assessed using a TapeStation D1000 System (Agilent, Santa Clara, CA, USA). All RNA samples used for sequencing had RIN values > 8.0.

RNA sequencing was performed by the University of Michigan sequencing core using an HiSeq 4000 system (Illumina, San Diego, CA, USA) and TruSeq reagents (Illumina) with 50bp single end reads as described (Gumucio *et al.*, 2019; Disser *et al.*, 2019). A total of 1μg of RNA from five rats from each group was analyzed. Read quality was assessed and adapters trimmed using fastp (Chen *et al.*, 2018). Based on fastp quality analysis, two samples from control group, one from the 7 day GSK2894631A 2mg group, and two from the 7 day 6mg GSK2894631A group were removed from further analysis. Reads were then mapped to the rat genome version RN6 and reads in exons were counted against RN6 Ensembl release 95 with STAR Aligner (Dobin *et al.*, 2013). Differential gene expression analysis was performed in R using edgeR (Robinson *et al.*, 2010). Genes with low expression levels (< 3 counts per million in at least one group) were filtered from all downstream analyses. A Benjamini-Hochberg false discovery rate (FDR) procedure was used to correct for multiple testing and FDR adjusted p values less than 0.05 were considered significant. Sequence data was deposited to NIH GEO (ascension number GSE130276).

For quantitative PCR (qPCR), RNA was first reverse transcribed into cDNA using iScript reagents (Bio-Rad, Hercules, CA, USA). qPCR was conducted in a CFX96 real time thermal cycler using SsoAdvanced SYBR green supermix reagents (BioRad). The 2^-ΔCt^ method was used to normalize the expression of mRNA transcripts to the stable housekeeping gene *Ppp1ca*. A listing of primer sequences is provided in Table 1.

**Table 1.**
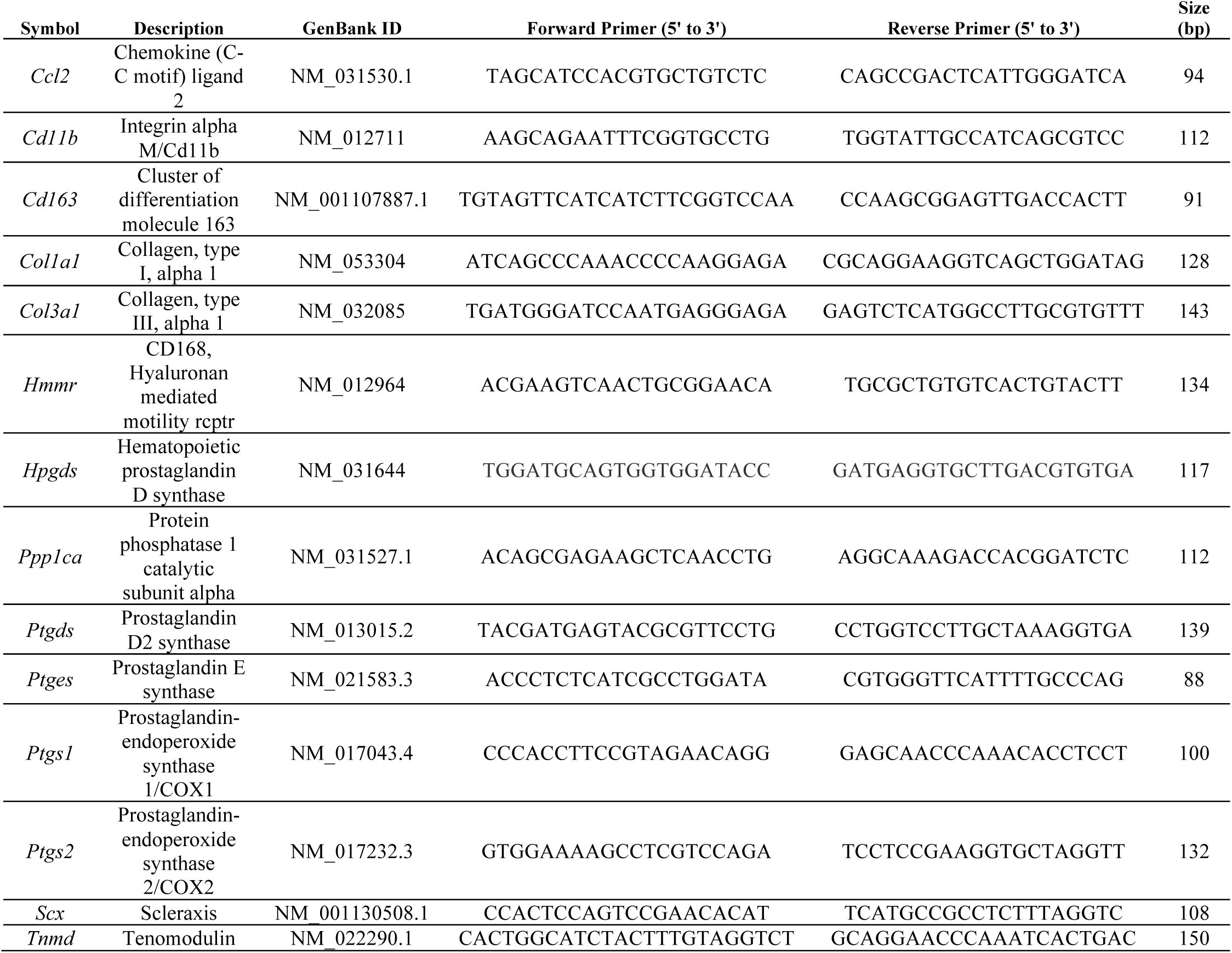
qPCR Primers. Sequences of primers used for qPCR.

### Statistics

Primary data was acquired in a blinded fashion. Values are presented as mean±SD. Statistical analyses of RNAseq data is described above. As the mechanical properties data in this study did not follow a Gaussian distribution, differences between groups were tested using a Kruskal-Wallis test followed by a Benjamini-Krieger-Yekutieli FDR correction (α=0.05) to adjust for multiple observations across groups. Gene expression, as measured by qPCR, was assessed using a Brown-Forsythe test followed by a Benjamini-Krieger-Yekutieli FDR correction (α=0.05). These analyses allowed for the assessment of differences between all treatment groups and control tendons, as well as differences within a time point and within a treatment dose. Prism (version 8.0, GraphPad, La Jolla, CA, USA) was used to perform statistical calculations.

## Results

An overview of the surgical procedure and study groups is shown in Figure 2A-B. All rats tolerated the surgical procedure, gavage, and drug treatment well, and there were no differences in body mass at the time of harvest (Figure 3A). As expected, the tenotomy and repair procedure resulted in inflammation and scar tissue formation, in particular around the areas of suture placement (Figures 2C-I). This resulted in an approximate 6-fold increase in the nominal cross-sectional area (CSA) of tendons across all repaired groups (Figure 3B). While the tendons became enlarged, no apparent gross differences in histological features were noted between the three treatment groups at either the 7 or 21 day time points (Figures 2D-I).

**Figure 2.**
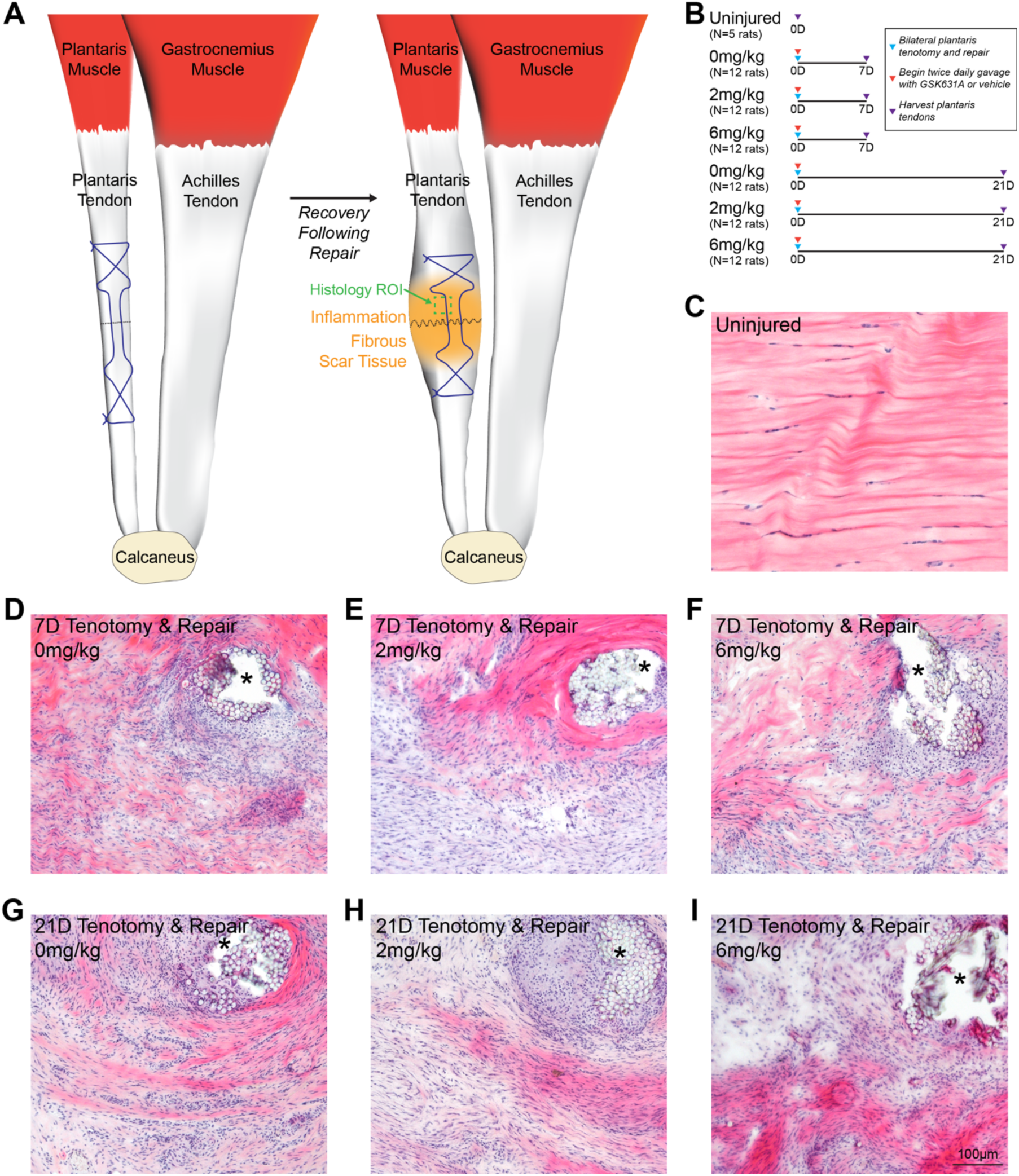
Overview of acute tenotomy and repair procedure, and representative histology of repaired plantaris tendons. (A) Overview of the surgical procedure, demonstrating a tenotomy (dashed black line) and Bunnell repair technique (suture pattern shown in blue) of the plantaris tendon. After the animals recover, inflammation and fibrous scar tissue will accumulate in the area of injury. The representative region of interest (ROI) for histology panels (C-I) is shown in green. (B) Overview of the study design and groups. (C-I) Hematoxylin and eosin histology stained sections from the midsubstance of plantaris tendons from (C) uninjured rats, and from rats treated with 0, 2, or 6mg/kg of GSK2894631A taken either 7 days (D-F) or 21 days (G-I) after acute tenotomy and repair. Areas of suture or suture resorption are shown with an asterisk. Scale bar for all histological sections is 100μm.

**Figure 3.**
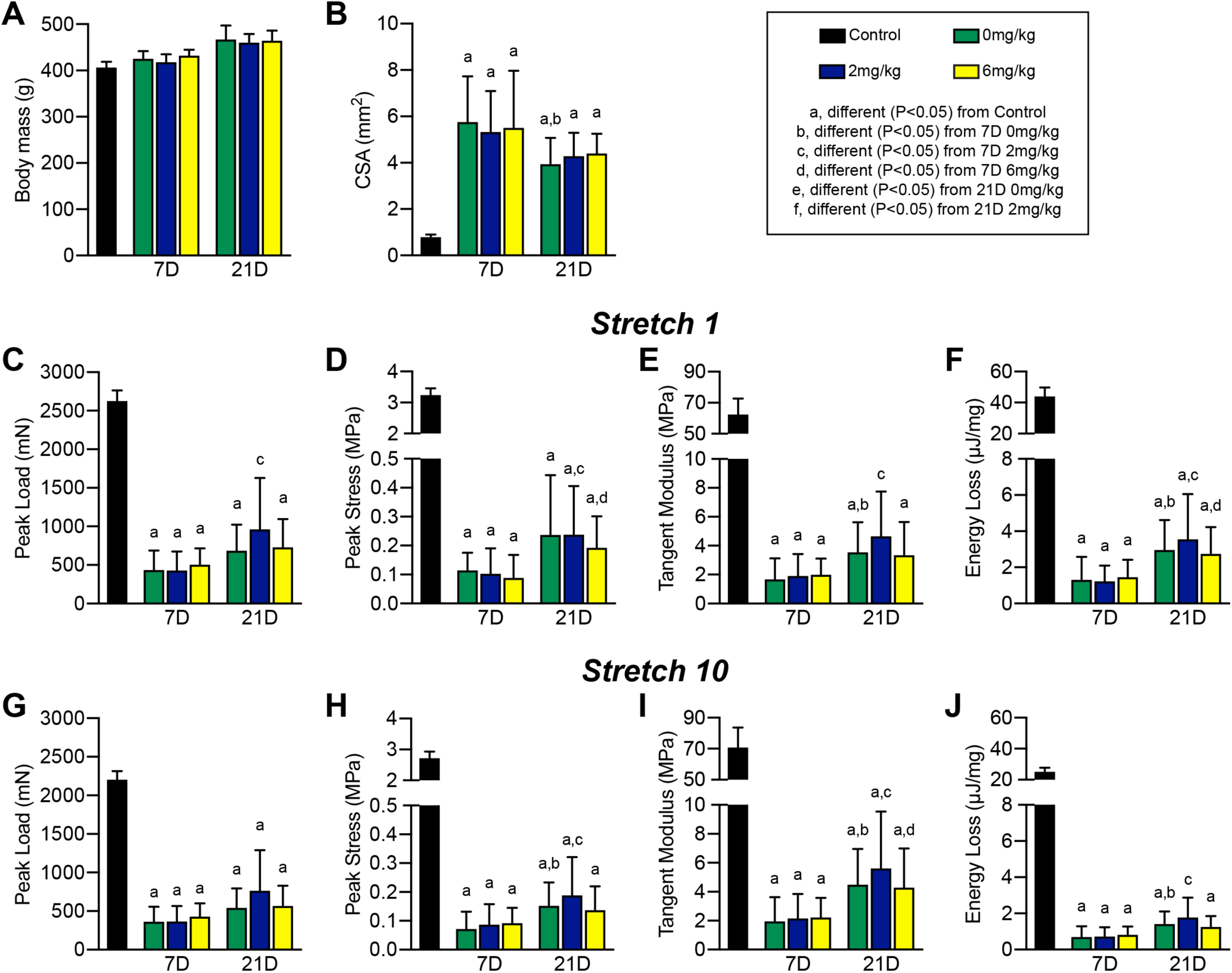
Mechanical properties of repaired plantaris tendons. (A) Animal body mass at the time of sacrifice, and (B) nominal cross-sectional area (CSA) of plantaris tendons. (C) Peak load, (D) peak stress, (E) tangent modulus, and (F) energy loss of tendons from the first of ten stretch cycles. (G) Peak load, (H) peak stress, (I) tangent modulus, and (J) energy loss of tendons from the last of ten stretch cycles. Values presented as mean±SD. Differences between groups were assessed using a Kruskal-Wallis test followed by a Benjamini-Krieger-Yekutieli FDR correction (α=0.05) to identify post-hoc differences between groups: a, different (FDR-adjusted P<0.05) from control tendons; b, different (FDR-adjusted P<0.05) from 7D 0mg/kg; c, different (FDR-adjusted P<0.05) from 7D 2mg/kg; d, different (FDR-adjusted P<0.05) from 7D 6mg/kg; e, different (FDR-adjusted P<0.05) from 21D 0mg/kg; f, different (FDR-adjusted P<0.05) from 21D 2mg/kg. N=5 tendons for controls, and N=12 tendons for each surgical repair group.

Mechanical properties testing was used to assess the functional impact of HPGDS inhibition on tendon repair, shown in Figures 3C-J. Tendons were stretched for 10 cycles with a total displacement of 10% original length (L_o_), and destructive testing was not performed to allow tendons to be preserved for histology. Broadly comparing control tendons to all repaired groups, peak load values were reduced by about 76% (Figure 3C and 3G), which is consistent with the observed disruptions to collagen fibrils in repaired tendons (Figures 2C-I). Peak stress was also lower in repaired groups by nearly 95% compared to uninjured tendons (Figures 3D and 3H), which is due to the reduction in peak load and the increase in CSA in repaired tendons (Figures 3B, 3C, and 3G). Tangent modulus and energy loss had similar reductions (Figures 3E, 3F, 3I, and 3J), likely due to an accumulation of fibrotic scar tissue (Figures 2C-I).

Comparing within repaired tendon treatment groups, the nominal cross-sectional area (CSA) of tendons across the 21D time point were about 24% lower than the 7D group (Figure 3B). There were no differences across time between the CSA of the three drug treatment groups, except for the 21D 0mg/kg group which was 32% smaller than 7D 0mg/kg tendons (Figure 3B). No differences in peak load at cycle 1 was observed across groups within a time point, although the 21D 2mg/kg group was about 2-fold higher than the 7D 2mg/kg group (Figure 3C). For peak stress, the 21D 0mg/kg and 21D 2mg/kg groups were about 2.3-fold higher than the corresponding 7D groups (Figure 3D). Tangent modulus and energy loss were not different between groups at a given time point, but for tangent modulus was 2-fold higher for the 21D 0mg/kg and 2mg/kg groups than they were at 7D, and for energy loss was 2.3-fold higher in all 21D groups compared to 7D tendons (Figures 3E-F). The results for changes in peak load, peak stress, tangent modulus, and energy loss at stretch 10 were generally similar to observations at stretch 1 (Figures 3C-J). Although the mechanical properties of repaired tendons across time points and treatment groups were inferior to uninjured tendons, the general shape of the stress-strain relationship remained similar (Figures 4A-C), and maintained a smooth morphology throughout the stretches indicating a relatively stiff repair callous. The loss in force over 10 stretch cycles was also generally similar between control tendons (Figure 4D), and in the treatment groups at the 7D and 21D time points (Figures 4E-F).

**Figure 4.**
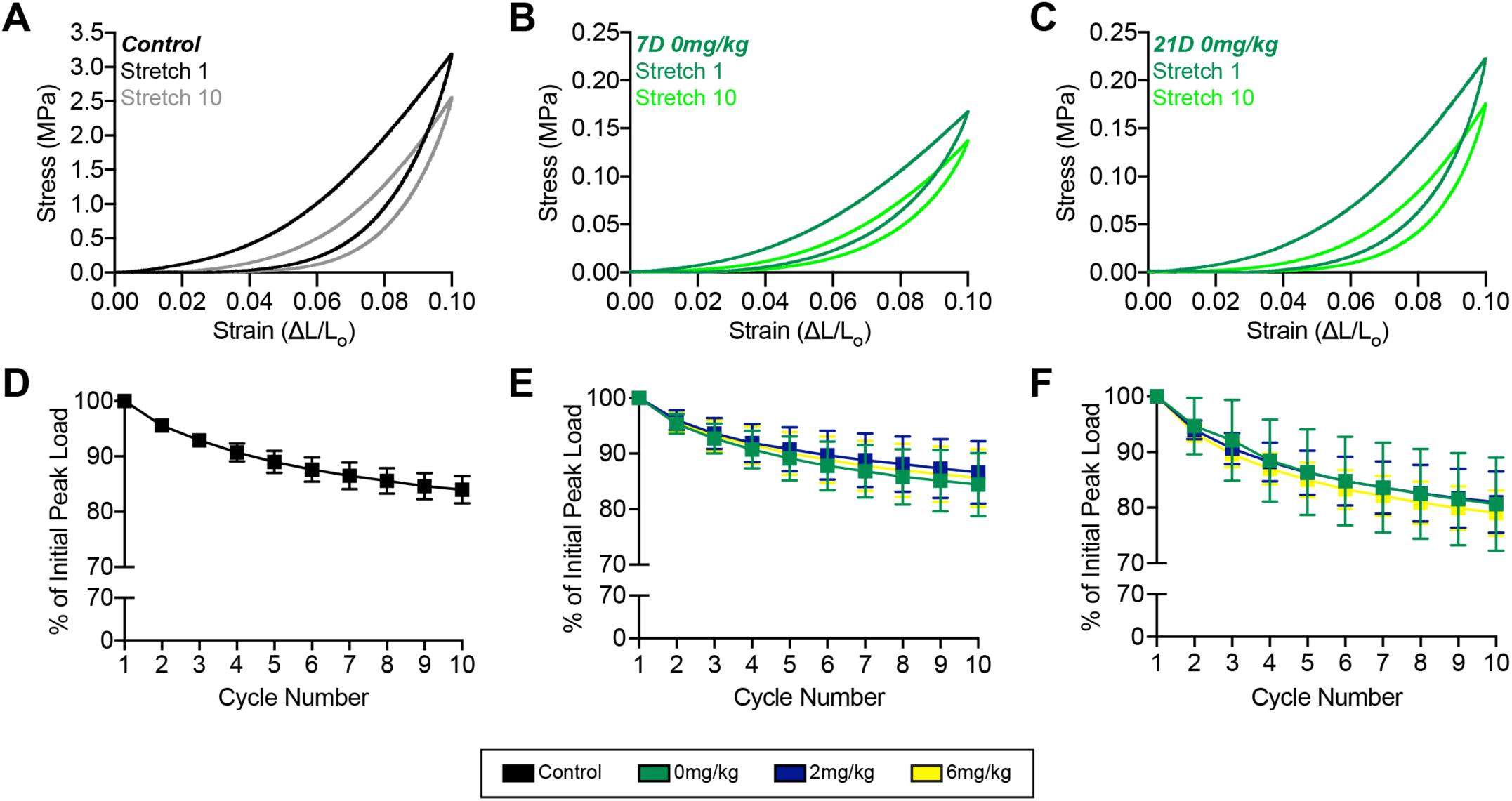
Stress-strain curves and peak load changes during stretch. Representative stress-strain response of a (A) control tendon, and (B) 7D 0mg/kg GSK2894631A (C) 21D 0mg/kg GSK2894631A repaired tendons from cycles 1 (darker color) and 10 (lighter color). Change in peak load across the ten cycles from (D) control tendons, and (E) 7D and (F) 21D repair groups.

We then performed RNA sequencing to comprehensively evaluate changes in transcript abundance. We first evaluated expression of genes involved in producing and sensing various prostaglandins in control and in 7D and 21D 0mg/kg groups. Plantaris tendons express *Ptgs1* and *Ptgs2*, which convert arachidonic acid (AA) into PGH_2_, in control and injured tendons (Figure 5). Tendons also robustly express enzymes which convert PGH_2_ into either PGE_2_ or PGF_2α_, as well as the receptors to sense these prostaglandins (Figure 5). However for PGD_2_, *Hpgds* was expressed at a low level and *Ptgds* was not detectable, nor were the PGD_2_ receptors *Ptgdr1* and *Ptgdr2* (Figure 5). *Ptgis* which converts PGH_2_ into PGI_2_ was not expressed in tendons, although the receptor *Ptgir* was expressed (Figure 5).

**Figure 5.**
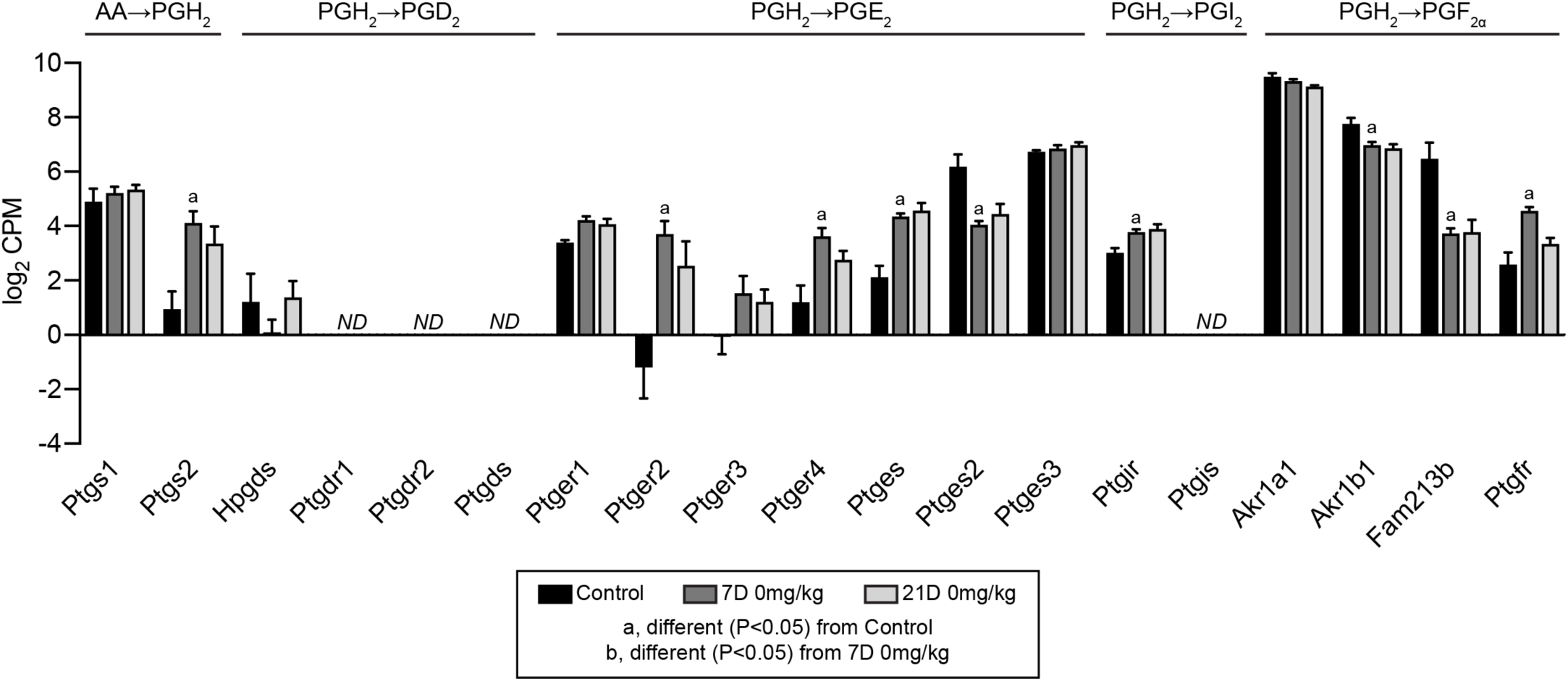
Prostaglandin synthesis RNAseq data. Expression in log_2_ counts per million mapped reads (CPM) for transcripts involved in the conversion of arachadonic acid (AA) to prostaglandin H_2_ (PGH_2_), and those which are involved in the conversion of PGH_2_ into PGD_2_, PGI_2_, and PGF_2α_, as well as the receptors for these prostaglandins. Values presented as mean±SD. Differences between groups tested with a FDR-adjusted t-test: a, different (FDR-adjusted P<0.05) from control tendons; b, different (FDR-adjusted P<0.05) from 7D 0mg/kg. N=3-5 tendons per group. ND, transcript not detected.

Finally, we analyzed global changes in RNAseq values. There were 3484 transcripts that had a FDR-adjusted P-value less than 0.05 (-log_10_P greater than 1.3) and were at least 1.5-fold upregulated (log_2_ fold change greater than 0.584) in 7D 0mg/kg tendons compared to controls, and 3222 transcripts that were significantly different (-log_10_P greater than 1.3) and were at least 1.5-fold downregulated (log_2_ fold change less than −0.584) in the 7D 0mg/kg group with respect to the control group (Figure 6A). By 21 days, only 82 transcripts were significantly upregulated and 43 were significantly downregulated compared to controls (Figure 6B). We then selected transcripts related to tendon healing and inflammation for further analysis across treatment groups and time points. Overall there appeared to be an effect of time since repair but not GSK2894631A treatment on regulating gene expression. For immune cell markers, compared to control tendons there was a general increase in the myeloid cell marker *Itgax*, the macrophage recruitment gene *Ccl2*, the pan-macrophage marker *Adgre1*, M1 macrophage markers *Ccr7* and *Cd68*, T cell markers *Cd3e* and *Cd8*, and the B cell marker *Ptprc* at 7 days, but the M2 macrophage markers *Cd163, Hmmr* and *Mrc1* were not different (Figure 6C). *Ptgs2* which is involved in the synthesis of PGH_2_ and *Ptges* which catalyzes PGH_2_ into PGE_2_ were upregulated, while another PGE_2_ synthesis enzyme *Ptges2* was generally downregulated 7 days after injury (Figure 6D). The ECM genes *Col4a1, Col6a1, Col12a*, and *Col14a1, Tnc* and *Vcan* were upregulated in 7D 0mg/kg tendons compared to controls, while *Col3a1* and the proteoglycans *Bgn* and *Fmod* were induced across treatment groups at 7 days (Figure 6E). *Mmp13* was upregulated in all 7 day groups, as was *Mmp14* which was also upregulated in the 21D 0mg/kg and 2mg/kg groups (Figure 6E). The growth factors *Igf1* and *Tgfb1* were upregulated in some of the 7 day groups compared to control tendons, as was the pro-inflammatory cytokine *Il1b* (Figure 6F). The early tenogenesis marker *Egr2* was generally upregulated at 7 days, while *Scx* was not different at any time point, and late tenogenesis markers *Mkx* and *Tnmd* were generally downregulated 7 days after injury (Figure 6G). Additionally, the myofibroblast marker *Acta2* and the tenocyte progenitor cell marker *Mcam* were upregulated in 7D 0mg/kg tendons (Figure 6G). We also performed qPCR to analyze select genes from injured tendons, and similar to RNAseq we generally observed very few differences between treatment groups at given time points (Table 2).

**Figure 6.**
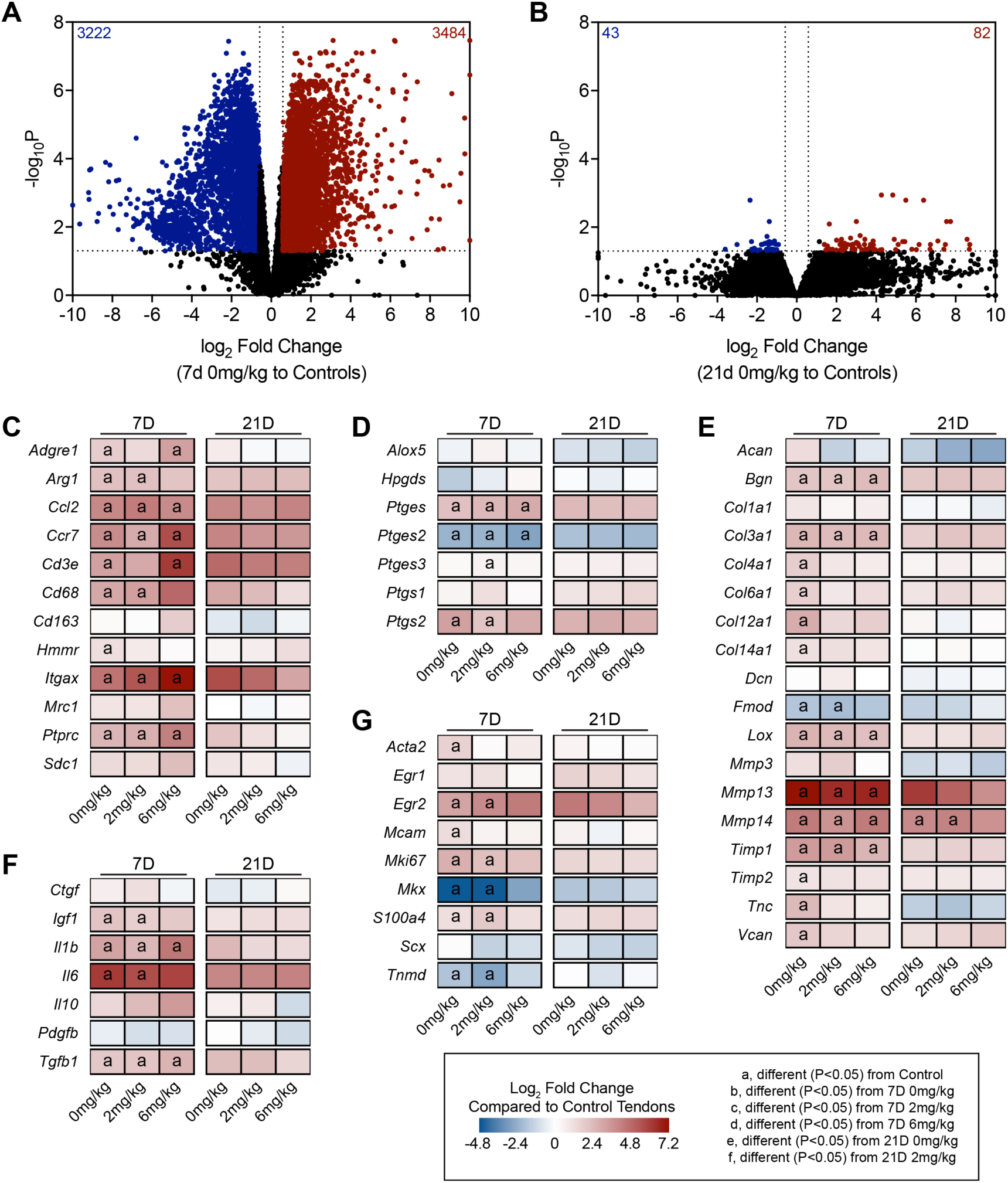
Overall RNAseq data. Volcano plots demonstrating log_2_ fold change and - log10 FDR-adjusted P-values of transcripts in the (A) 7D 0mg/kg GSK2894631A and (B) 21D 0mg/kg GSK2894631A groups, compared to control tendons. Heatmaps demonstrating expression of selected transcripts that are (C) inflammatory cells markers, (D) prostanoid metabolism genes, (E) involved in ECM synthesis and remodeling, (F) growth factors and cytokines, and (G) markers of tenogenesis. Data are log_2_ fold change in expression of each treatment group normalized to control tendons. Differences between groups tested with a FDR-adjusted t-test: a, different (FDR-adjusted P<0.05) from control tendons; b, different (FDR-adjusted P<0.05) from 7D 0mg/kg; c, different (FDR-adjusted P<0.05) from 7D 2mg/kg; d, different (FDR-adjusted P<0.05) from 7D 6mg/kg; e, different (FDR-adjusted P<0.05) from 21D 0mg/kg; f, different (FDR-adjusted P<0.05) from 21D 2mg/kg. N=3-5 tendons per group.

**Table 2.**
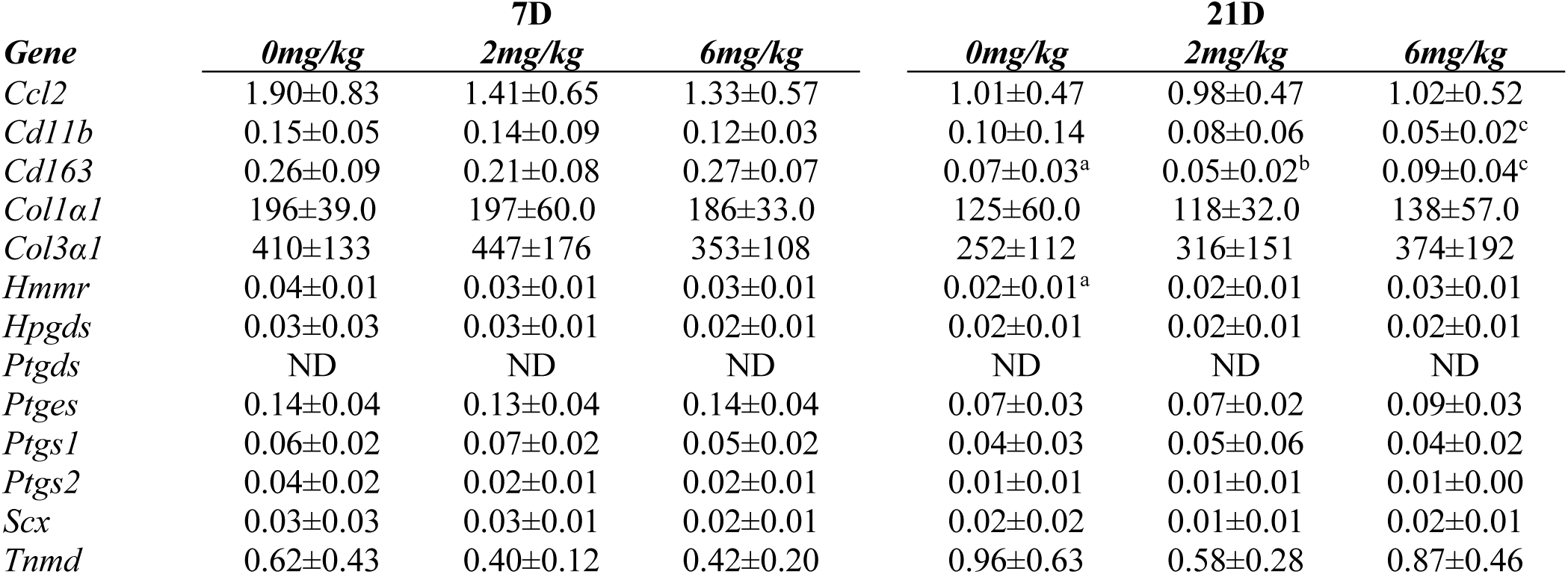
qPCR. Gene expression in injured tendons. Target genes are normalized to the stable housekeeping gene *Ppp1ca*. Values presented as mean±SD. Differences between groups were assessed using a Brown-Forsythe test followed by a Benjamini-Krieger-Yekutieli FDR correction (α=0.05) to identify post-hoc differences between groups: a, different (FDR-adjusted P<0.05) from 7D 0mg/kg; b, different (FDR-adjusted P<0.05) from 7D 2mg/kg; c, different (FDR-adjusted P<0.05) from 7D 6mg/kg; d, different (FDR-adjusted P<0.05) from 21D 0mg/kg; e, different (FDR-adjusted P<0.05) from 21D 2mg/kg. N=6 tendons per group. ND, not detected.

| *Gene* | **7D** |  |  | **21D** |  |  |
| --- | --- | --- | --- | --- | --- | --- |
|  | *0mg/kg* | *2mg/kg* | *6mg/kg* | *0mg/kg* | *2mg/kg* | *6mg/kg* |
| *Ccl2* | 1.90±0.83 | 1.41±0.65 | 1.33±0.57 | 1.01±0.47 | 0.98±0.47 | 1.02±0.52 |
| *Cd11b* | 0.15±0.05 | 0.14±0.09 | 0.12±0.03 | 0.10±0.14 | 0.08±0.06 | 0.05±0.02<sup>c</sup> |
| *Cd163* | 0.26±0.09 | 0.21±0.08 | 0.27±0.07 | 0.07±0.03<sup>a</sup> | 0.05±0.02<sup>b</sup> | 0.09±0.04<sup>c</sup> |
| *Col1a1* | 196±39.0 | 197±60.0 | 186±33.0 | 125±60.0 | 118±32.0 | 138±57.0 |
| *Col3a1* | 410±133 | 447±176 | 353±108 | 252±112 | 316±151 | 374±192 |
| *Hmmr* | 0.04±0.01 | 0.03±0.01 | 0.03±0.01 | 0.02±0.01<sup>a</sup> | 0.02±0.01 | 0.03±0.01 |
| *Hpgds* | 0.03±0.03 | 0.03±0.01 | 0.02±0.01 | 0.02±0.01 | 0.02±0.01 | 0.02±0.01 |
| *Ptgds* | ND | ND | ND | ND | ND | ND |
| *Ptges* | 0.14±0.04 | 0.13±0.04 | 0.14±0.04 | 0.07±0.03 | 0.07±0.02 | 0.09±0.03 |
| *Ptgs1* | 0.06±0.02 | 0.07±0.02 | 0.05±0.02 | 0.04±0.03 | 0.05±0.06 | 0.04±0.02 |
| *Ptgs2* | 0.04±0.02 | 0.02±0.01 | 0.02±0.01 | 0.01±0.01 | 0.01±0.01 | 0.01±0.00 |
| *Scx* | 0.03±0.03 | 0.03±0.01 | 0.02±0.01 | 0.02±0.02 | 0.01±0.01 | 0.02±0.01 |
| *Tnmd* | 0.62±0.43 | 0.40±0.12 | 0.42±0.20 | 0.96±0.63 | 0.58±0.28 | 0.87±0.46 |

## Discussion

Tendon tears in adult animals heal through the formation of a fibrovascular scar, with inferior mechanical properties that disrupt proper force transmission, limit performance, and increase the susceptibility for a reinjury (Yang *et al.*, 2013; Nourissat *et al.*, 2015; Freedman *et al.*, 2017). Inflammation is a hallmark of tendon tears, and various prostaglandins are produced throughout the stages of tendon injury and repair (Su & O’Connor, 2013). PGD_2_ plays a role in promoting inflammation in various diseases, including skeletal muscle and nerve injury, and the inhibition of PGD_2_ production has produced promising results in animal models and early clinical trials (Thurairatnam, 2012; Santus & Radovanovic, 2016). Given these encouraging findings, we tested the hypothesis that a potent and specific inhibitor of PGD_2_ synthesis, GSK2894631A, would improve the recovery of tendons following an acute injury and repair. Although the test compound was well tolerated, and a handful of genes were differentially regulated across treatment groups, the targeted inhibition of PGD_2_ did not impact the functional repair of tendons after injury.

NSAIDs and coxibs, which inhibit the production of PGH_2_ from arachidonic acid, are used to treat pain and inflammation after tendon injury. However, many studies have shown that the use of these drugs reduces or delays tendon healing (Ferry *et al.*, 2007; Dimmen *et al.*, 2009; Hammerman *et al.*, 2015), which is similar to observations in other musculoskeletal tissues (Cohen *et al.*, 2006; Su & O’Connor, 2013; Dueweke *et al.*, 2017; Lisowska *et al.*, 2018). PGH_2_ is metabolized by specific synthases to produce other prostaglandins, such as PGD_2_, PGE_2_, PGF_2α_, and PGI_2_, that modulate inflammation (Trappe & Liu, 2013). PGD_2_ plays an important role in promoting inflammation, and inhibiting the HPGDS and PTGDS enzymes which produce PGD_2_ from PGH_2_ generally results in favorable clinical outcomes (Thurairatnam, 2012; Santus & Radovanovic, 2016).

In the current study we found that inhibiting HPGDS had no appreciable effect on tendon healing. HPGDS is expressed in various immune cells, such as Th2 lymphocytes, antigen-presenting cells, macrophages, mast cells, megakaryocytes, and eosinophils (Thurairatnam, 2012; Kern *et al.*, 2017), and while little is known about the adaptive immune response in tendon, macrophages are known to accumulate after tendon injury (Marsolais *et al.*, 2001; Sugg *et al.*, 2014). We evaluated the expression of several markers of macrophages and adaptive immune cells, and although we generally observed an upregulation in these markers after injury, HPGDS was detected at a low level in tendon tissue and was surprisingly downregulated in most groups after injury, while other enzymes involved with prostaglandin synthesis, such as PTGES and PTGS2, were upregulated in injured tendons. The two receptors for PGD_2_, PTGDR1 and PTGDR2, were also not detected in any tendon samples. There was no clear pattern for the effect of HPGDS inhibitor treatment on growth factors, cytokines, ECM components or tenocyte markers. Combined, these results suggest that GSK2894631A does not impact tendon healing in a positive or negative manner, likely due to an absence of PGD_2_ producing enzymes and PGD_2_ receptors in healing tendon tissue.

There are several limitations to this work. We only evaluated two time points, chosen to be representative of the late inflammatory phase (7 days) and well into the proliferative and regenerative phases of tendon healing (21 days), and it is possible that PGD_2_ producing enzymes are expressed later and have a role in modulating late stages of tendon healing. It is also possible that PGD_2_ producing enzymes are expressed earlier in the repair process, but even if they are, any effects that would have occurred early on would not seem to have any impact on functional healing at later stages. Only male rats were evaluated in this study, as tendon ruptures occur three times more frequently in men than women (Ganestam *et al.*, 2016), however we think the results are likely applicable to both males and females. We measured transcriptional changes with RNAseq and qPCR but did not measure proteomic changes in tendons, and changes in transcript levels may not reflect changes in protein abundance. Finally while we analyzed PGD_2_ biology in plantaris tendons of rats, it is possible that other tendons, or even different species or strains of rats, do express HPGDS at a higher level, and that there could be a therapeutic role for a PGD_2_ inhibitor in these instances.

## Conclusion

In the current study, based on exciting reports from other tissues and conditions, we tested the hypothesis that the targeted inhibition of HPGDS would improve tendon healing following an acute plantaris tenotomy and repair. The findings of this study have lead us to reject this hypothesis, as inhibiting PGD_2_ did not affect tendon healing, likely due to the low abundance of HPGDS after injury. Although this is a negative finding, we still think this can inform the potential clinical use of PGD_2_ inhibitors. While we used an acute injury model in this study, chronic tendon tears often result in substantial muscle atrophy (Davis *et al.*, 2015; Davies *et al.*, 2015), and there is compelling data that inhibiting PGD_2_ can improve the recovery of skeletal muscle after injury and protect against atrophy (Mohri *et al.*, 2009; Thurairatnam, 2012). Therefore blocking PGD_2_ production in a way that improves muscle healing without impacting tendon could be a substantial advance from the current clinically available prostaglandin synthase inhibitors, NSAIDs and coxibs, which generally delay healing and result in inferior functional outcomes for both muscle and tendon tissue.

## Acknowledgements

This study was funded by GlaxoSmithKline. KBS was supported by a fellowship from the NIH (F32-AR067086). Bioinformatics support was provided by a grant from the Tow Foundation for the David Z Rosensweig Genomics Center at the Hospital for Special Surgery. The authors wish to acknowledge the assistance of Mr. Patrick Stillson and Dr. James Markworth with animal procedures.

## Author Contributions

DCS, KBS, HFK, ACH, and CLM designed research; DCS, KBS, JRT, JBS, DO, and CLM performed research; DCS, KBS, JRT, JBS, DO, and CLM analyzed data; HFK and ACH contributed critical reagents; DCS, KBS, DO, and CLM wrote the paper. All authors reviewed and approved the final version of the manuscript.

